# Kainic Acid Pig Model of Hippocampal Epilepsy

**DOI:** 10.1101/2025.01.22.634312

**Authors:** Filip Mivalt, Daniela Maltais, Inyong Kim, Jiwon Kim, Patrik Began, Andrea Duque Lopez, Veronika Krakorova, Bailey Winter, Cheng Yen Kuo, Shelja Sharma, Elizabeth S. Harty, Luke H. Kim, Nicholas Gregg, Dan Montonye, Christopher Gow, Kai Miller, Jamie Van Gompel, Kent Leyde, Vaclav Kremen, Su-youne Chang, Gregory A. Worrell

## Abstract

Translational large-animal models that can accommodate human-scale implantable devices are essential for advancing neuromodulation therapies in epilepsy. This study establishes a kainic acid (KA)-induced porcine model of mesial temporal lobe epilepsy (mTLE) using clinical imaging, stereotactic surgery, and a fully implantable neural stimulation and recording (INSR) device designed for humans.

Seven pigs (six KA-treated and one saline control) underwent MRI-guided stereotactic implantation of electrodes targeting bilateral hippocampus (HPC) and anterior thalamus (ANT), followed by intra-hippocampal KA or saline infusion. Local field potentials (LFP) were recorded continuously with synchronized video monitoring. Seizures and LFP interictal epileptiform-like discharges (IEDs) were quantified using validated automated detectors. Histology was performed in the saline control and the longest surviving KA-treated pig.

Intra-hippocampal KA infusion induced acute status epilepticus in all (6/6) treated pigs. Four animals survived to chronic monitoring with spontaneous seizures observed in three pigs (2,733 seizures; mean duration of 27.2 ± 17.6 seconds). IEDs were observed in bilateral HPC of all animals, including saline control, with higher rates in the lesioned HPC (p <0.0001). While the IED morphology is consistent with epileptiform activity; IEDs alone are not specific for epilepsy and physiological transients (e.g. sharp-wave ripples) and injury-related hyperexcitability or strain-specific hyperexcitability cannot be excluded. Histological analysis revealed patchy neuronal loss and cytoarchitectural changes in HPC.

This porcine model reproduces electrophysiological features of human mTLE. This approach provides a powerful translational bridge for developing and testing next-generation INSR and neuromodulation strategies in freely behaving large animals.

## Introduction

Epilepsy is a prevalent neurological disorder affecting approximately 50 - 60 million people worldwide [1]. Beyond recurrent seizures, epilepsy is often associated with psychiatric comorbidities including anxiety, depression [2]–[4], memory impairments [3], cognitive deficits [2], [3], and sleep disturbances [5]–[7], reflecting complex network pathophysiology [2]. These comorbidities are thought to arise from circuit reorganization [8], neuroinflammation [9], neurodegeneration [10], [11], and functional network abnormalities [12]. Mesial temporal lobe epilepsy (mTLE) with hippocampal seizures exemplifies the challenges of drug-resistant epilepsy and is a compelling target for advanced neuromodulation [13]–[22].

Electrical brain stimulation is an established therapy for drug-resistant epilepsy, reducing seizure burden with the anterior nucleus of the thalamus (ANT) and hippocampus (HPC) being common neuromodulation targets [18], [21]. However, complete seizure freedom is rare, and stimulation parameter optimization is a lengthy process taking months to years [18], [19], [21]. The extended parameter optimization timeline is impacted by the sporadic occurrence of seizures as the main outcome measure, limited-duration local field potential (LFP) recordings and inaccurate patient-reported seizure counts [18], [19], [21]–[23]. These challenges motivate the development of next-generation implantable neural stimulation and recording (INSR) devices [13], [16].

Developing and testing next-generation neuromodulation devices for epilepsy can be accelerated by appropriate animal models that (i) accommodate human-scale devices, (ii) replicate human electrophysiology, behavior, and (iii) demonstrate predictive validity for therapeutic responses. Rodent models have been extensively used in epilepsy research [24]; however, their small size and limited neuroanatomical complexity constrain their utility for device development. Large animals, such as dogs [25], [26] and pigs [27]–[29], provide a gyrencephalic brain structure and sufficient size to support multimodal imaging, surgical procedures, and implantable devices designed for human use.

Chemical induction of seizures with intra-hippocampal kainic acid (KA) has been widely applied in rodent models [24], [30] and recently demonstrated in a porcine model [28]. In rodents, localized intra-amygdala or intra-hippocampal KA infusion produces a period of acute status epilepticus, followed by a latent period and the emergence of chronic epileptiform activity characterized by interictal epileptiform-like transients (IEDs), high-frequency oscillations (HFO) and spontaneous seizures [30]–[35]. These models also reproduce key neuropathological features of human drug resistant mTLE [36], [37].

In contrast to rodent models, relatively few studies have explored large animal models of mTLE [24]. Zhu et al. [28] described a porcine model using intra-hippocampal KA injections, with intermittent (2-hour daily) electrophysiological recordings under light anesthesia to quantify IEDs, HFOs, and seizures. In that study, 66% of pigs receiving 10µL intra-hippocampal KA exhibited IEDs and 25% developed spontaneous seizures. However, the impact of light anesthesia during limited electrophysiology recordings on seizure counts remains uncertain, and histological characterization was not reported. Here, we extend this prior work [28] by investigating an intra-hippocampal KA porcine model of mTLE using a neurotechnology platform integrating advanced clinical neuroimaging protocols, image-guided stereotactic targeting, and an INSR device designed for human use. The approach enables chronic synchronized video and continuous local field potential (LFP) recordings in freely behaving large animals (Figure 1). The rechargeable INSR device enables continuous long-term LFP recording without the need for anesthesia. Furthermore, we perform histological analysis to assess hippocampal structural changes.

**Figure 1:**
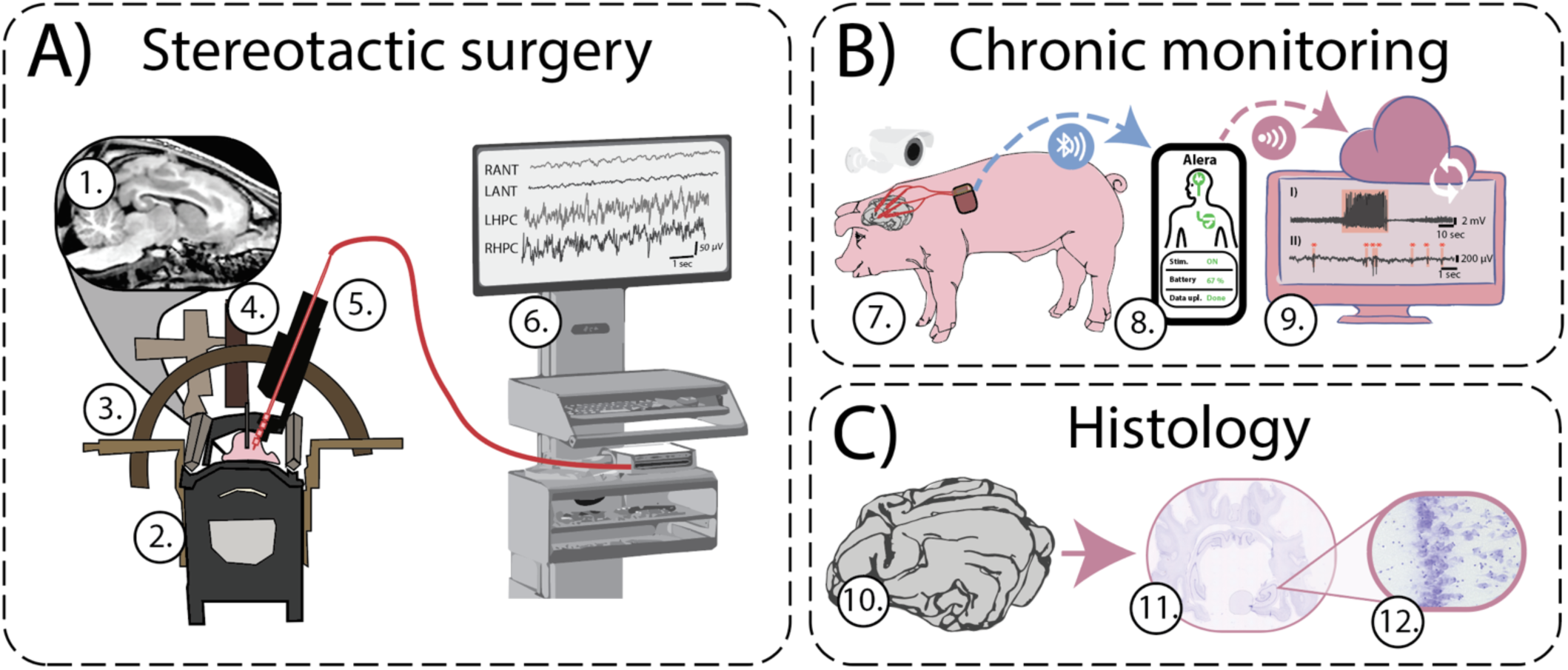
Neurotechnology for Large Animal Epilepsy Models. **A) Stereotactic implantation surgery** using 1) 3-Tesla pre-op MRI. 2) Custom MRI compatible stereotactic frame for head fixation. 3) Stereotactic Leksell frame compatible with Brainlab stereotactic planning software (Brainlab AG, Munich, Germany) for electrode targeting. 4) Microinjector for kainic acid (KA) infusion. 5) Electrode and cannula delivery system to implant electrodes and inject KA. 6) Intra-operative workstation for electrophysiology monitoring, single pulse electrical stimulation, and tracking the local field potential (LFP) changes induced by KA infusion. **B) Chronic electrophysiology monitoring** 7) Implanted animals were chronically monitored with synchronized video and LFP recordings using an implantable four lead neural stimulating and recording device that wirelessly transfers LFP recordings using a Low Energy Bluetooth 8) Android phone application for programming. 9) Cloud with automated LFP signal processing for seizure and IED detection. **C) Brain Histology** was performed to investigate structural changes in the 10) explanted brains for 11) gross structural and 12) microscopic histologic changes.

## Results

We developed a porcine model of mTLE using stereotactic intra-hippocampal KA injection combined with chronic electrophysiological and video recordings in freely behaving animals using a human-grade INSR.

### Subjects

Seven domestic Yorkshire pigs (Sus scrofa) were included: six received intra-hippocampal KA (S1-6) and one control animal received saline (S7). Animals were implanted with four leads (four contacts per lead) targeting bilateral HPC and ANT and monitored using synchronized video and continuous LFP recordings.

All procedures were conducted at Mayo Clinic and adhered to the National Institutes of Health Guidelines for Animal Research under (IACUC protocol A00006870) in accordance with NIH and ARRIVE guidelines.

### Acute Surgery and KA Induction of Status Epilepticus

Four leads were implanted in bilateral HPC and ANT using Magnetic Resonance Imaging (MRI) guided stereotactic surgery.

The intraopertaive LFPs demonstrated lower amplitudes and power spectral density in the ANT compared to the HPC (ANT: 0.80 ± 3.92 µV/Hz vs HPC: 8.77 ± 5.83 µV/Hz; p < 0.0001, Mann-Whitney U-test) consistent with reports from human recordings [14].

Single pulse evoked potentials (SPEPs) were recorded in two pigs (S5-S6) intraoperatively to evaluate Papez circuit functional connectivity [36], [37]. ANT→HPC responses exhibited longer latencies than HPC→ANT responses (16.64 ± 5.88 ms vs. 8.44 ± 2.25 ms, p <0.0001, Mann-Whitney U-test), consistent with asymmetric network functional connectivity. HPC →HPC SPEP demonstrated higher amplitude response compared to the ANT →HPC (304.81 ± 176.19 µV vs 64.02 ± 33.13 µV, p <0.0001, Mann-Whitney U-test) (Figure 2).

**Figure 2.**
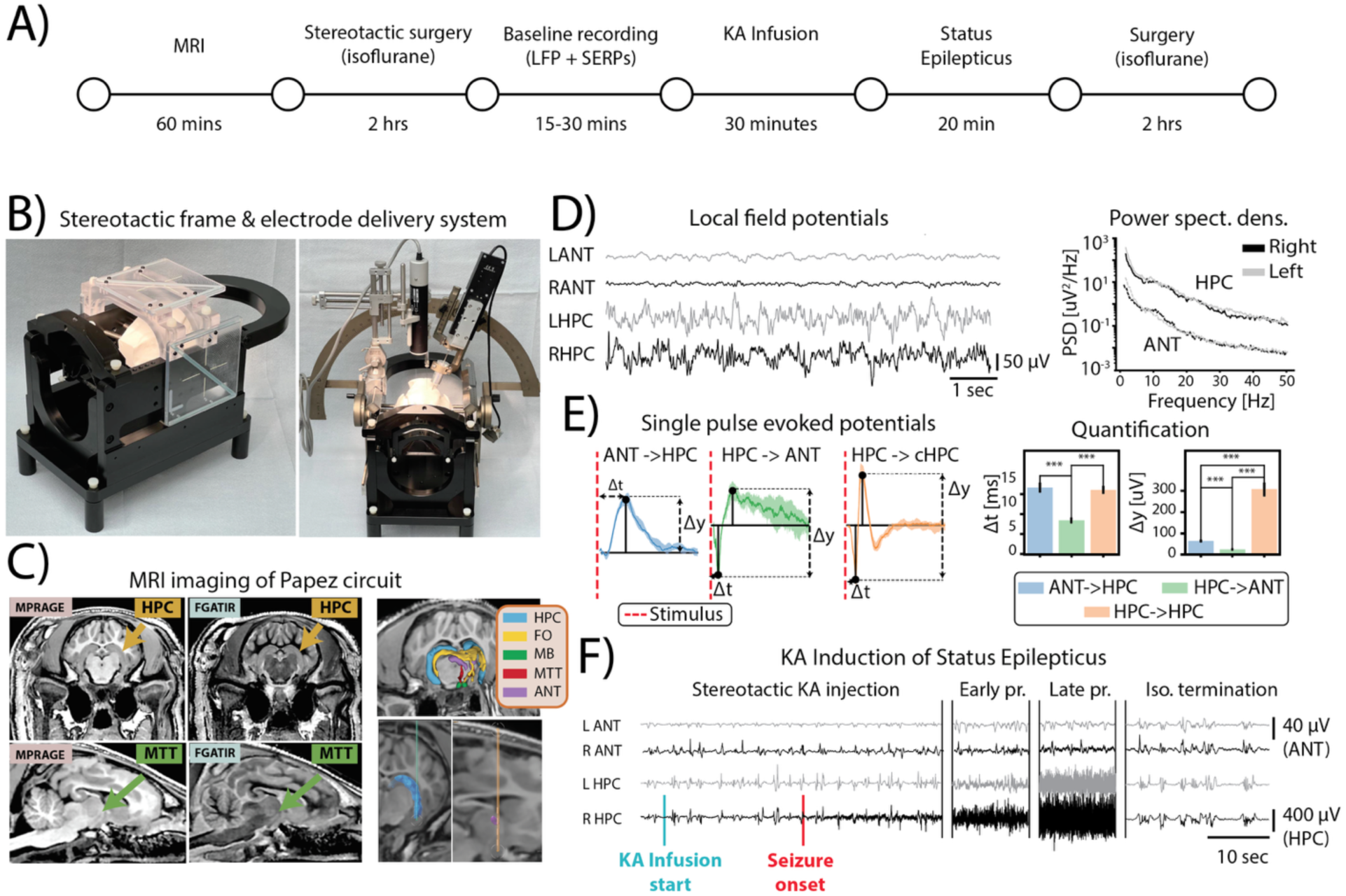
Intraoperative Electrophysiology of Papez Circuit and Acute Status Epilepticus. **A) Experimental Timeline. B) MRI compatible stereotactic frame.** Custom MRI compatible head frame with jaw and skull set screws for head fixation with N-bar fiducial box. Leksell arc-frame and computer-controlled drive for electrode and chemotoxin delivery. **C) MRI scans utilized for pre-operative planning.** MPRAGE and FGATIR were utilized for targeting and volumetric rendering of hippocampus (HPC), fornix (FO), mammillary bodies (MB), mammalothalamic track (MTT) and anterior nucleus thalamus (ANT). **D) Representative intra-operative local field potential** traces from bilateral ANT and HPC recording at baseline with corresponding power spectral density. **E)** Single Pulse Evoked Potentials from ANT to HPC (ANT→HPC), HPC to ANT (HPC→ANT) and HPC to contralateral HPC (HPC→cHPC). The latency of initial negative response peak (Δt) and peak to peak magnitude(Δy) were evaluated. **F) Kainic Acid (KA) Seizure Induction.** Intra-HPC **i**njection of KA provoked prolonged status epilepticus that was terminated with Isoflurane after 20 minutes. The onset of right HPC seizure after KA infusion. The seizure propagates into the left HPC and right ANT.

Intra-hippocampal KA infusion (1µg/µl; total volume 4 – 16 µl) induced focal electrographic seizures at the injection site, with propagation to the ANT and contralateral HPC (Figure 2). Seizures and prolonged SE were acutely observed intra-operatively in all animals except the saline control (S7).

The targeting accuracy was confirmed with post-operative Computed Tomography and histology.

### Recovery and Acute Seizure Dynamics

In KA-treated pigs during the first hour upon emergence from anesthesia, bilateral hippocampal seizure activity persisted and occasionally progressed to focal to bilateral tonic-clonic seizures (FBTCS). These seizures included tonic extension of a forelimb, lateral head deviation, and generalized clonic movements. Limb movements resembling “running” were also intermittently observed. While these overt behavioral manifestations were primarily confined to the first hour, electrographic seizure activity in the HPC persisted for up to 12 hours post-operatively. Electrographic seizure patterns were characterized by evolving rhythmic high-frequency activity at onset followed by increased amplitude and spatial spread.

Subjects S1 and S6 were euthanized within 12 hours of surgery. Subject S1 received the highest KA dose (16 µl KA) and had refractory SE that was unresponsive to intravenous diazepam. Continued electrographic seizures were recorded post-operatively with the INSR device despite intravenous diazepam and the animal was euthanized per our protocol. Subject S6 (6µl KA) had prolonged post-operative seizures and received intravenous diazepam (0.5 - 1 mg/kg). The pig experienced cardiopulmonary arrest, and resuscitation was unsuccessful. Autopsy revealed cardiac pathology, including dilated ventricles with thin myocardial walls, possibly a pre-existing condition contributing to cardiopulmonary arrest and death. KA-pigs S3 and S4 were euthanized on day 10 and 19, respectively after surgery due to wound infections around the INSR device per the IACUC protocol for humane euthanasia in setting of complications.

The control animal (S7) did not develop acute seizure or status epilepticus.

### Automated Epileptiform Seizure and IED Detection

Automated IED and seizure detectors previously developed for humans and canines [13] were validated in pigs against manual annotations by a human expert (Figure 3A). The automated IED detection algorithm achieved a sensitivity of 0.87 and a Positive Predictive Value (PPV) of 0.68 when compared to the expert annotations. The automated seizure detector achieved a Receiver Operating Characteristic Area Under the Curve (AUC) of 0.93 and a Precision-Recall Curve AUC of 0.84 when compared to the expert annotations (Figure 3A).

**Figure 3.**
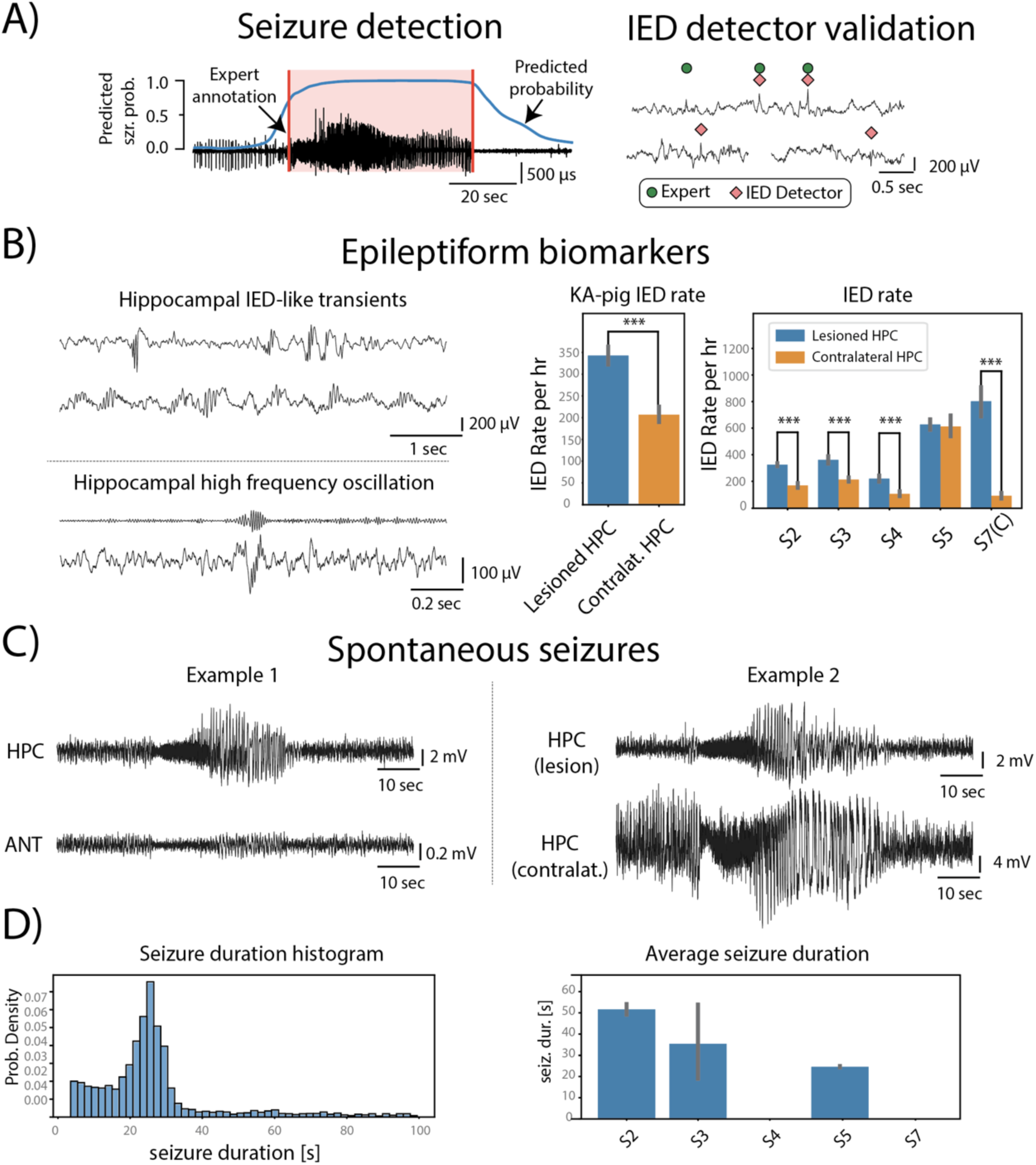
Interictal Epileptiform-like Discharges (IED) and Seizures. **A) Automated seizure and IED detection.** *Left)* Representative seizure detection with the algorithm predicted probability (blue line) overlaid on the local field potential (LFP) and expert annotation (red shaded). *Right)* Representative IED detections (red diamonds) and expert annotations (green circles). **B) IEDs during chronic monitoring.** *Left)* Representative IEDs and high frequency oscillations. *Right)* IED rates in the lesioned and contralateral hippocampus (HPC). The IED rates are higher in the lesioned HPC (KA & control saline) compared to the contralateral HPC. **C) Spontaneous Seizures during chronic monitoring.** *Left)* A representative right HPC seizure recorded in right HPC and right anterior nucleus of thalamus (ANT). *Right)* HPC seizure with propagation to contralateral HPC. **D)** Histogram of automatically detected seizure durations. Average seizure duration for each subject. Note, S4 and the control S7 did not have any seizure detections.

### Chronic Monitoring, Seizures and IEDs in behaving pig mTLE model

The four KA-treated surviving pigs (S2, 3, 4 and 5) and the control pig (S7) were maintained and monitored using video and LFP recordings for a total of 433 days (S2: 271 days, S3: 10 days, S4: 19 days, S5: 86 days, S7-control: 47 days) (Table 1.).

**Table 1:**
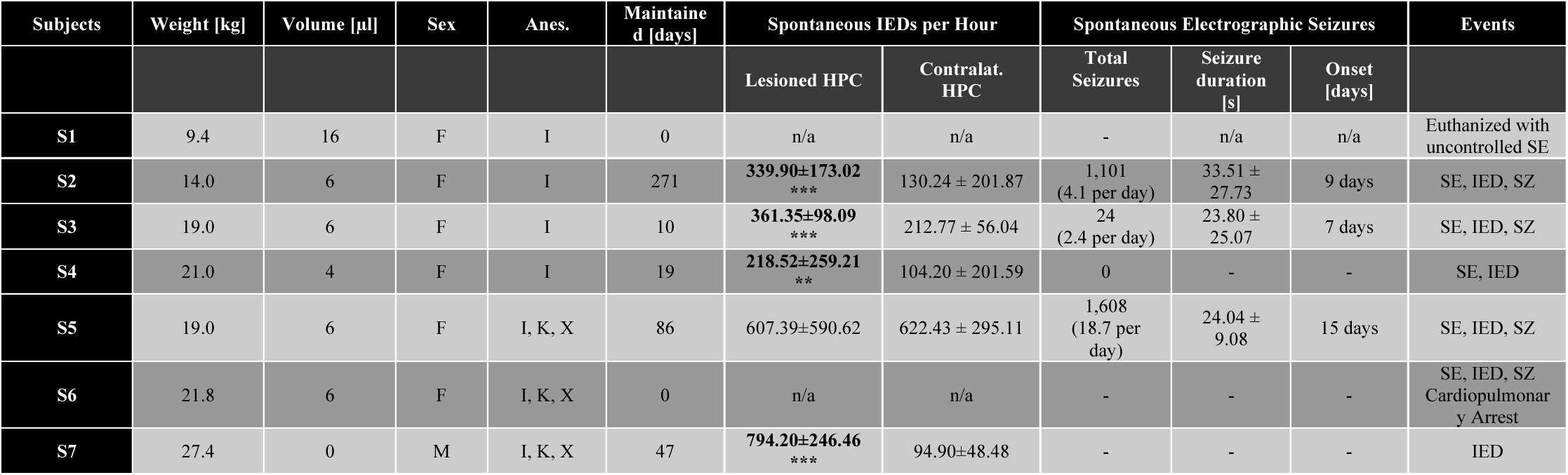
Summary of Animal Outcomes and Electrophysiology. Seven domestic Yorkshire pigs, six females (F) and one male (M) were used for intrahippocampal Kainic Acid (KA; 1 µg/µl) model. S7 was a control pig. The weight at the time of implant; total injected KA dose; hippocampal Interictal epileptiform discharges (IEDs), seizures (SZ), and status epilepticus (SE) (* - p<0.05; ** - p<0.01, *** p<0.001, Mann-Whitney – lesioned HPC vs Contralateral HPC). Isoflurane (I), Ketamine (K), Xylazine (X).

| Subjects | Weight [kg] | Volume [µl] | Sex | Anes. | Maintained [days] | Spontaneous IEDs per Hour |  | Spontaneous Electrographic Seizures |  |  | Events |
| --- | --- | --- | --- | --- | --- | --- | --- | --- | --- | --- | --- |
|  |  |  |  |  |  | Lesioned HPC | Contralat. HPC | Total Seizures | Seizure duration [s] | Onset [days] |  |
| S1 | 9.4 | 16 | F | I | 0 | n/a | n/a | - | n/a | n/a | Euthanized with uncontrolled SE |
| S2 | 14.0 | 6 | F | I | 271 | 339.90±173.02<br>*** | 130.24 ± 201.87 | 1,101<br>(4.1 per day) | 33.51 ± 27.73 | 9 days | SE, IED, SZ |
| S3 | 19.0 | 6 | F | I | 10 | 361.35±98.09<br>*** | 212.77 ± 56.04 | 24<br>(2.4 per day) | 23.80 ± 25.07 | 7 days | SE, IED, SZ |
| S4 | 21.0 | 4 | F | I | 19 | 218.52±259.21<br>** | 104.20 ± 201.59 | 0 | - | - | SE, IED |
| S5 | 19.0 | 6 | F | I, K, X | 86 | 607.39±590.62 | 622.43 ± 295.11 | 1,608<br>(18.7 per day) | 24.04 ± 9.08 | 15 days | SE, IED, SZ |
| S6 | 21.8 | 6 | F | I, K, X | 0 | n/a | n/a | - | - | - | SE, IED, SZ<br>Cardiopulmonary Arrest |
| S7 | 27.4 | 0 | M | I, K, X | 47 | 794.20±246.46<br>*** | 94.90±48.48 | - | - | - | IED |

IED transients were observed in all surviving subjects. The IED morphologies included spike and wave discharges, high frequency ripple oscillations [38] and polyspikes, all of which have been reported in previous KA seizure models [28], [31], [39] (Figure 3). We observed independent and synchronous bilateral HPC IEDs with higher hourly IED rates in the lesioned HPC (lesioned 352.68 ± 338.49 vs. contralateral 174.59 ± 269.71, p <0.0001, Mann-Whitney U-test) in all subjects including the control pig. These findings are insufficient to infer causality, but are consistent with that electrode implantation and injection alone inducing network hyperexcitability independent of epilepsy.

Three out of four of the KA-treated pigs had spontaneous seizures (S2, S3, S5) occurring an average of 10.33 ± 3.40 days post-KA infusion. A total of 2,733 electrographic seizures (mean seizures/day S2: 4.1, S3: 2.4, S4: 0, S5: 28.7) were recorded (Table 1). Mean seizure duration was 27.16 ± 17.62 seconds. (Table 1). On visual review of synchronized video and LFPs, we observed multiple behavioral seizure correlates including brief behavioral arrest, generalized tonic-clonic convulsive seizures and myoclonic jerks.

The control animal (S7) did not have electrographic seizures during surgery or over the 47-day chronic monitoring period.

### Histology

Quantitative histology was performed on the control pig (S7) and the KA-treated pig (S5). The histology of the KA-treated HPC revealed visually apparent pathological changes in the hippocampal subfields (CA1, CA3, and dentate gyrus (DG)) including patchy pyramidal and granule cell loss, and the presence of pyknotic nuclei [40] (Figure 4). The pathologic changes for each structure specifically included (Figure 4.):

**Figure 4.**
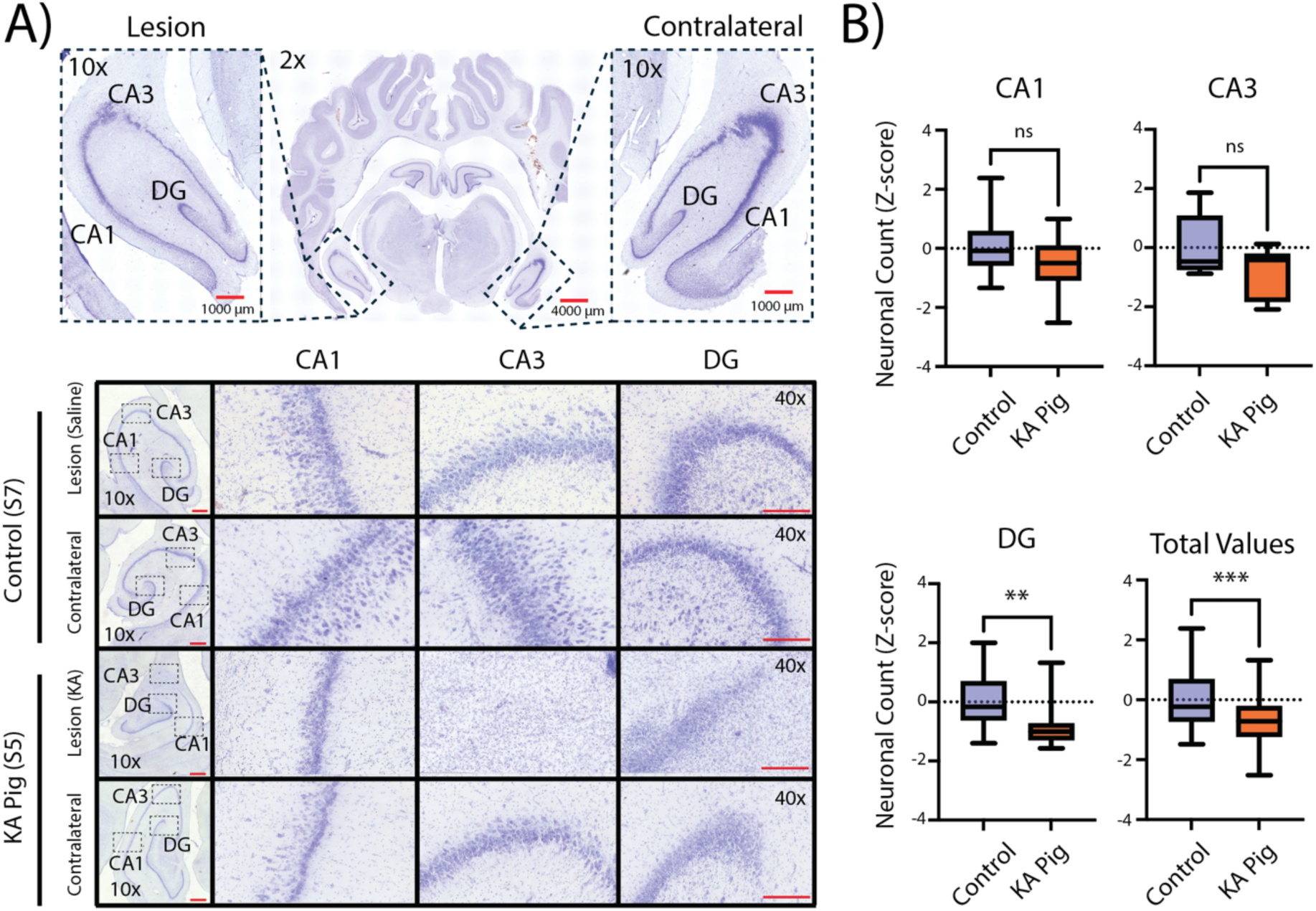
Histology from the Lesioned and Contralateral Hippocampus. **A) Coronal brain section with cresyl violet staining.** *Top)* Representative image at low (2x) magnification of the lesioned and contralateral hemispheres. Boxed areas (CA1, CA3, and Dentate (DG) at 10x magnification). *Bottom)* Representative examples from control and chronic pig (10x & 40x magnification). In the KA lesioned hippocampus patches of pyramidal cell loss were observed in CA3 and CA1 region and granule cell loss in the DG. **B)** Comparisons of normalized neuronal cell counts between the saline injection control animal and the KA injection lesioned pig. The DG showed a marked loss of granular cells in the KA pig. The aggregated cell counts across HPC subfields were reduced in the lesioned HPC. Hippocampal regions. CA1: Cornu Ammonis 1; DG: Dentate Gyrus; CA3: Cornu Ammonis 3; Hi: Hilus. Scale bars: 4000 μm at 2x, 1000 μm at 10x, 200 μm at 40x.

CA1: Observed patches of pyramidal cell loss and presence of dark pyknotic nuclei. Quantitative analysis of normalized pyramidal cell counts showed a non-significant trend towards pyramidal cell loss (p = 0.145).

CA3: Observed patches of pyramidal cell loss, where certain ROIs showed a complete absence of pyramidal neurons. Quantitative analysis of normalized pyramidal cell counts showed a non-significant trend towards pyramidal cell loss (p= 0.293)

DG: Observed patches of granule cell loss and presence of dark pyknotic nuclei, which corresponded to a significant reduction in granule cell counts (p= 0.003) (Figure 5B, right).

**Figure 5:**
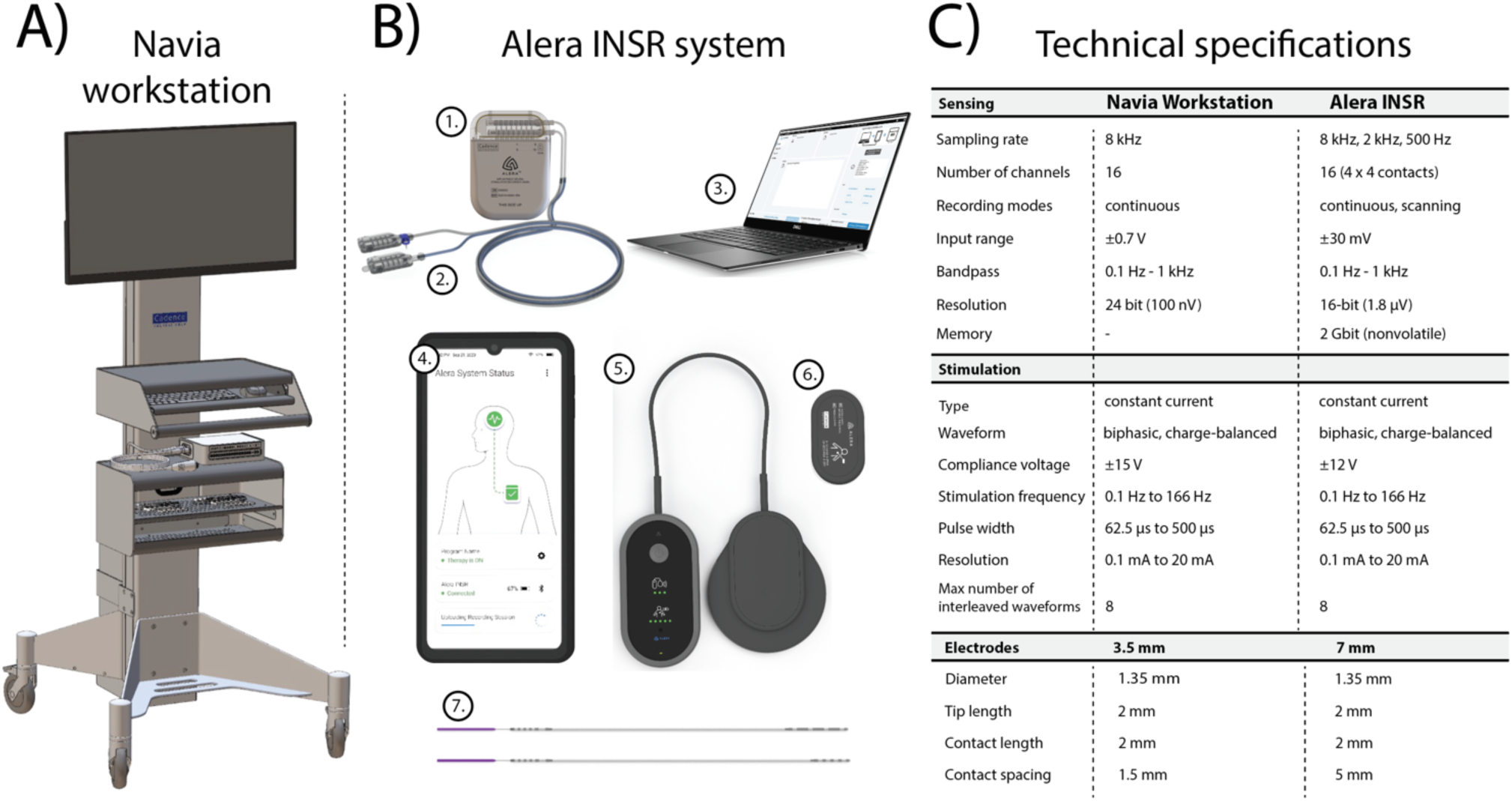
Local Field Potential (LFP) Recording Instrumentation. **A) Navia Workstation (NW)** is a 16-channel LFP recording and electrical stimulation platform. The NW enables high fidelity LFP sensing and recording (0.1 – 1000 Hz: Sampling 8 kHz) and programmable electrical stimulation. **B) Alera Implantable Neural Stimulation and Recording (INSR) system**. The INSR is a rechargeable, 16-channel sensing (0.1 – 1000 Hz: Sampling up to 8 kHz, 2 Gb memory), stimulation and recording device. The Alera system includes: 1) 16 channel INSR supporting four leads (4-contacts each) with 2) electrode extensions with skull mounted bifurcation port, 3) programmer (Tablet computer), 4) controller (Android mobile phone), 5) charger, 6) control magnet and 7) electrode leads with 3.5 mm and 7 mm span. The INSR wirelessly communicates with the controller, relaying information such as the INSR battery level, operational status, and recording status. The controller stores recorded LFP data downloaded from the INSR. The programmer is used to generate stimulation therapy programs that are uploaded to the INSR via the controller. **C) Detailed device specifications**.

Neuronal counts aggregated over all the examined subfields demonstrated a significant overall decrease in the KA-treated HPC (p = 0.001)

## Discussion

In this study, we developed a porcine intra-hippocampal KA model of mTLE and performed chronic electrophysiological recordings, concurrent video monitoring and post-mortem HPC histology. The model demonstrated several key structural and electrophysiological features of human mTLE, supporting its use as a translational model. Large animal models of epilepsy represent a vital step in the development and testing of human use INSR devices.

Our study builds upon the prior work of Zhu et al. [28], who established the feasibility of the porcine intra-hippocampal KA model with intermittent electrophysiological recordings under light anesthesia. The primary novelty of our work is the development of an integrated platform for clinical-grade imaging, acute intra-operative recording with SERP-based probing of Papez circuit and chronic electrophysiology in behaving animals. Key distinctions include the implementation of a chronic, fully implantable, human-grade INSR device, the specific targeting and functional characterization of the Papez circuit (ANT-HPC, HPC-HPC) using SPEPs.

Additionally, 24/7 synchronized video-LFP monitoring (1 second synchronization accuracy) in freely behaving animals. The substantially higher electrographic seizure counts captured in our cohort (2,733 seizures in 3 behaving animals during a total of 386 days) compared to the prior study (46 seizures in 4 animals recorded for 2 hours per day under light anesthesia for 28 days) [28] is likely attributable to our continuous LFP monitoring without anesthesia and reporting of all electrographic seizure events with a subtle or no behavioral correlates.

### KA Dose and Seizure Outcomes

We investigated a range of intra-hippocampal KA doses (4µl, 6µl, and 16µl) to establish a porcine mTLE model. All doses successfully induced acute status epilepticus (SE); however, the 16µl dose resulted in refractory SE that required euthanasia. The 6µl dose, used in most pigs (4/6), successfully induced chronic IED and spontaneous seizures in surviving animals, demonstrating its effectiveness in producing the desired mTLE phenotype. While this dose group (6 µl KA) did experience mortality, the cause were attributed to non-dose-related factors, including post-operative infection and likely pre-existing cardiac pathology. Therefore, while our findings show that a 6µl dose is effective, future studies with a larger cohort are required to fully optimize the protocol for consistent long-term epilepsy outcomes, and determining evidence for drug resistant and neuromodulation efficacy.

### Seizure and IED Detector Validation

The IED and seizure detection algorithms were previously validated in both human and canines [13], [44]. Here the IED detectors yielded good performance in the pig model without further hyperparameter tuning. We acknowledge the sample size is a limitation and an area for more extensive validation in future work.

The emergence of IED transients following implantation of the HPC depth electrodes in all animals, including control animal, suggests that electrode implantation can produce injury-related hyperexcitability. Furthermore, the increased IEDs in the saline-infusion control compared to the contralateral HPC suggests that the electrode implantation procedure itself and saline injection may contribute to network hyperexcitability, although causality cannot be determined in this study. Notably, IEDs alone are not specific for epilepsy and physiological transients and injury-related hyperexcitability or strain-specific hyperexcitability cannot be excluded [42]. The electrode leads (1.3 mm diameter) used in this study are designed for human use and are relatively large for the pig HPC and produce tissue trauma. The human electrode lead implant and 6 µl saline injection is likely not an entirely benign event. Notably, the saline control animal had IEDs but did not have seizures.

A key methodological distinction that likely explains the higher seizure counts observed in our porcine KA model is the use of continuous, long-duration electrographic monitoring with inclusion of all electrographic seizure events, rather than reliance on intermittent sampling or behaviorally anchored seizure detection. In rodent KA models, even when telemetry is employed, seizure burden is often underestimated because intermittent recordings or behavioral scoring can miss subclinical or nonconvulsive electrographic seizures [43]. Indeed, Dudek and colleagues emphasize that seizures with minimal or absent behavioral correlates are difficult to detect with video monitoring alone, highlighting the necessity of continuous EEG to capture the full spectrum of epileptiform activity [43]. Similarly, studies using chronic radiotelemetry demonstrate that virtually continuous (24 h/day) recordings over months reveal progressive increases in seizure frequency, clustering, and a predominance of nonconvulsive electrographic events [44]. These findings underscore that seizure counts are highly dependent on sampling strategy: continuous monitoring detects not only overt convulsive seizures but also frequent electrographic events that can dominate overall seizure burden. Thus, the higher seizure rates in our pig model likely reflect a more complete accounting of epileptic activity enabled by uninterrupted recording and rigorous automated detection of all electrographic seizures, rather than a fundamental biological discrepancy between species.

### Behavioral Seizure Correlates

Our characterization of behavioral seizures focused primarily on overt events with clear motor manifestations, such as focal to bilateral tonic-clonic seizures that included jerking and falling, or sudden awakenings from sleep. A detailed classification of seizures impacting behavior was beyond the scope of this initial study. Therefore, a systematic assessment for seizures with more subtle behavioral correlates, such as those limited to brief behavioral arrest, was not performed. However, we did note subtle behavioral changes, such as a sudden cessation of activity, which post-hoc inspection of the LFP confirmed were correlated with hippocampal electrographic seizures. Developing and implementing specific behavioral assays to formally assess functionally impairing seizures is a key objective for future studies.

### Hippocampal Histology

The quantitative HPC histology performed on the KA-pig shows patches of pyramidal cell loss in CA1 and CA3, but only a trend in cell loss when averaged over the individual subfields. The DG, however showed significant overall cell loss. It is important to note that human mTLE with HPC seizures has varied histology with 25% (38/150) of mTLE cases coming to surgery having normal histology [45]. Whereas, human mTLE related to mesial temporal sclerosis shows significant pathology within CA1, CA3 and DG [46]. Our findings in the pig KA-model fall between these extremes found in human mTLE.

### Comparison with human mTLE

Similar to rodents the intrahippocampal-KA pig model demonstrates a relatively short latent period for the emergence of spontaneous seizures. In this study a latent period was observed, with spontaneous seizures beginning an average of 10.33 ± 3.40 days post-insult, similar to previous porcine model report [28] is also similar to rodent studies, e.g. ∼ 1 - 2 weeks for mice and rats [32], and shorter than the months or years often described in human acquired epilepsy [8], [47]–[49]. The accelerated timeline in animal models is likely a feature of the direct intra-hippocampal KA infusion, which serves as a potent and targeted insult, potentially speeding up the network reorganization that leads to chronic epilepsy [30]. Future studies could explore lower KA doses to investigate if a more prolonged latent period can be established, which may also more closely model the human condition.

### Implications for Neuromodulation Device Development

The porcine mTLE model provides a unique opportunity to advance neuromodulation therapies agnostic to the device. The relatively large gyrencephalic brain and ability to accommodate multiple electrode leads and INSR devices designed for humans should be useful for testing neuromodulation and sensing paradigms [13], [14], [17], [23], [50]–[54].

### Limitations and Future Directions

This study is limited by the small sample size, which may affect the generalizability of findings. Most notable is the single control animal available in this study. Importantly, the control animal did have IED-like transients, but did not have intra-operative status epilepticus or chronic spontaneous seizures supporting the epileptogenic nature of intra-hippocampal KA. The control pig did develop IED-like transients, which are a known non-specific signature of brain injury and not necessarily epilepsy [55], [56]. We were also unable to assess post-KA SPEP changes intra-operatively due to persistent focal status epilepticus masking any evoked activity. This leaves a gap in understanding the effect of KA on acute evoked responses.

The quantitative histological analysis is limited to a single control and single chronic KA-HPC pig. This is inadequate to formulate a complete picture of the pathological changes of intra-HPC KA induced epilepsy. However, quantitative cell count analysis comparing the KA- and Saline-control animals did confirm DG, but not in CA1 and CA3, cell loss. This may reflect methodological limitations, small sample size, sparse HPC histological sampling along the anterior-posterior axis, and secondary pathological changes related to electrode injury in the contralateral control side.

The range of clinically relevant information including if the animals are drug resistant, have progressive epilepsy, or comorbidities (such as memory and sleep disruption) cannot be ascertained from the current study.

Future studies should include a larger cohort to validate findings and further refine KA dosing for improved survival and chronic seizure outcomes. Lastly, incorporating behavioral assessments, such as responsiveness, memory, mood, and sleep alongside electrophysiological monitoring could enhance our understanding of seizure-related comorbidities in this model [57], [58]. Finally, exploring additional neuromodulation targets within the Papez circuit could provide insights into broader therapeutic applications for epilepsy-related cognitive and emotional deficits.

## Conclusion

We successfully demonstrated an intra-HPC KA model of mTLE in pigs. Using chronic electrophysiology with synchronized video monitoring we identified seizures and IED activity. This model demonstrates reliable induction of acute status epilepticus, following hippocampal KA injections, and in surviving animals chronic epilepsy with spontaneous IEDs and seizures. The model supports chronic monitoring in freely behaving animals and in the future can be used for testing novel sensing and stimulation paradigms. Our findings underscore the translational potential of a porcine mTLE model for addressing unmet clinical needs of novel stimulation therapies in drug-resistant epilepsy.

## Methods

### Animals

Animals were housed on a 12/12 light cycle with ad libitum access to water and once-daily feeding. Animals were housed in temperature-controlled rooms (45% humidity, 21°C temperature) with elevated floors that met all size, material and sanitation requirements according to Guide for the Care and Use of Laboratory Animals [59]. Animals were provided mats and daily enrichment via assorted treats, chew toys, and human interaction. Animals were socially housed, except during post-surgery period daily assessed by a team of veterinarians and technicians. All animals were humanely euthanized at the conclusion of the study using Phenobarbital.

### Pre- and Post-Surgical Imaging

Pre-operative planning was conducted using a 3.0 Tesla MRI (GE Medical Systems). Magnetization-prepared rapid gradient echo (MPRAGE) and fast gray matter acquisition T1 inversion recovery (FGATIR) sequences were selected for their enhanced gray-white matter contrast [60], [61]. These MRI sequences, commonly used for electrode lead placement planning in human deep brain stimulation (DBS), provided visualization of the mammillothalamic tract (MTT) for precise direct ANT targeting [62]. Post-operative electrode placement was confirmed through computed tomography (CT) (Somatom Flash, Siemens Inc.), co-registered with the pre-operative MRI. Images were analyzed in Brainlab (Brainlab AG, Munich, Germany) and 3D Slicer [63] (Figure 2).

### Stereotactic & Recording Instrumentation

A custom stereotactic surgery platform consisting of an MRI compatible, machined plastic head fixation frame (Supplement) and a modified Leksell arc-frame (Elekta Inc., Stockholm, Sweden) compatible with a stereotactic targeting software Brainlab (Brainlab AG, Munich, Germany) was used for electrode implantation and KA infusion. The stereotactic frame was designed to accommodate up to 40 kg pigs, with 15 kg – 25 kg being an optimal weight. See supplementary material.

A 16-channel Cadence Navia Workstation (Cadence Neuroscience Inc., Seattle, WA) LFP sensing (0.1 Hz - 1 kHz bandpass, 8 kHz sampling rate, 24-bit resolution,) and programmable stimulation was used for acute intraoperative electrophysiology monitoring.

Chronic LFP monitoring was conducted using a fully implantable, 16-channel Cadence Alera INSR device (Cadence Neuroscience Inc., Seattle, WA). This system records from two digitally selected bipolar derivations (0.1 Hz - 1 kHz bandpass, 8 kHz sampling rate, 16-bit resolution and a 2 Gbit nonvolatile memory). The INSR system includes the implantable device, a handheld programmer/controller for wireless bi-directional data transfer via Bluetooth Low Energy, and a charger (Figure 5).

### Stereotactic Surgery and Kainic Acid Infusion

Pre-operative MRI scans and stereotactic targeting software (Brainlab AG, Munich, Germany) and porcine brain atlas [64] were used for segmentation and targeting of HPC and ANT, both common targets in mTLE neuromodulation therapy [13], [16], [18], [19], [21], [22]. Four 4-electrode leads were implanted through skull burr holes, secured with “dog-bone” anchors and bone screws.

Anesthesia was induced and maintained with isoflurane for the stereotactic surgery. The initial four pigs (S1-S4) underwent KA infusion under continuous low-level isoflurane and the last two pigs (S5, S6) with a combined ketamine-xylazine anesthesia protocol following a 30-minute isoflurane washout (Table 1). This anesthesia change was implemented because of concern that isoflurane, a GABA-A agonist and direct kainate receptor antagonist, anti-convulsant properties could pharmacologically confound the acute KA status epilepticus [30], [65].

One pig (S7) underwent an identical stereotactic surgery and intra-hippocampal infusion with an equivalent volume (6 µl) of sterile saline instead of KA as a control for the effects of the hippocampal injection procedure itself.

At surgery the externalized electrode leads were connected to the bedside electrophysiology workstation for acute monitoring using sterile electrode extensions. KA in six pigs and sterile saline in the control pig were stereotactically delivered to the hippocampus along a separate track using a Hamilton syringe positioned 1-2 mm from the HPC recording electrode contact. KA (1µg/µl) or saline was administered in 2 µL increments at a rate of 1 µl per minute, with a 5-minute interval between doses. We investigated 16 µl, 6 µl, and 4 µl KA total dose and 6 µl saline dose in the control pig. Focal hippocampal status was terminated with isoflurane anesthesia after 20 minutes. Intravenous benzodiazepines (0.5 – 1.0 mg/kg) were utilized on as needed basis during surgery and recovery to suppress clinical seizures.

After conclusion of the acute recording, the four electrode leads were tunneled under the skin and connected to bifurcation ports mounted on the skull and two-wire extensions (8-contact extensions) were tunneled to the implantable neural sensing and recording (INSR) device (Cadence Neuroscience Inc., Seattle, WA) positioned behind the right scapula. Post-surgical CT scans were used to confirm electrode placement.

### LFP and Video Recording and Analysis

Cadence Navia workstation was utilized to record LFPs and probe the Papez circuit using SPEP. The LFP referential recordings from the ANT and HPC using the 16 electrode contacts (4 contacts on each of the 4 implanted leads) were obtained during surgery before (Figure 2C) and with intrahippocampal KA infusion (Figure 2E). The SPEP were measured as a response to a single pulse stimulation with 6 mA current, 250µs pulse width at 0.1 Hz. A visual review was utilized to intra-operatively identify seizures.

INSR devices were implanted behind the right scapula and used for chronic LFP recording with a range of sampling rates (500 Hz, 2 kHz or 8 kHz). The sampling rate of 500 Hz was utilized for chronic LFP recordings. Higher sampling rates were utilized to record short LFP recordings of 1 minute to 10 minutes periodically. The INSR enables stimulation from the 16 contacts and LFP recording from two digitally selected bipolar electrode derivations (Figure 5).

Bipolar LFP channel recordings were derived from the raw acute LFP traces recorded using the Navia workstation for each brain location. The workstation LFPs were subsequently filtered using anti-aliasing filter and downsampled to 500 Hz for further analysis. A narrow-band notch filter with quality factor 10 was utilized to suppress 60 Hz powerline noise in the workstation LFP data. Downsampled workstation LFP data and longitudinal INSR LFP data were stored at 500 Hz sampling, and the same signal processing protocol was utilized for both workstation and INSR data.

LFP signals were filtered between 2 Hz and 100 Hz using a 101^st^-order finite response (FIR) filter. Previously developed Python libraries [13], [14], [57] were utilized to process LFP signals.

A USB 1080P Full HD Camera (SVPRO, Shenzhen, China) was used to collect video at 30 frames per second. Video was recorded using Blue Iris software (Perspective Software LLC, Northville, Michigan, USA). The camera was placed above the animal pen with a coverage of the pen area. The video stream was recorded with the computer clock timestamp. The internal clock of the INSR was synchronized to the computer clock with a 1 second precision before every initiation of a recording. LFP traces and simultaneously recorded video files were manually reviewed using Cyber PSG software (Certicon a.s., Prague, Czech Republic).

### Automated IED and Seizure Detection Algorithm Validation

An IED and seizure detector previously validated in humans and canines [13], [44] was validated for the pig study. A human expert (CYK) annotated IEDs and seizures from long-term recordings (222 IEDs and 42 seizures) for validation. All data annotated were utilized solely for testing of the previously developed detectors. In this study no additional training or tuning of detector hyperparameters was performed. After validation in the pig the IED and seizure detection algorithms were utilized to detect seizures and IEDs in the freely behaving pigs.

### Histology

Two of the seven pigs were used for histological evaluation: S5 and S7 (control). The explanted brains were fixed in 4% paraformaldehyde and stored in 30% sucrose in 1X Phosphate-Buffered Saline. Brains were frozen with dry ice and cut to 60-µm thick coronal sections on a sliding microtome (Leica Biosystems, Wetzlar, Germany) in serial order starting from the corpus callosum anteriorly to the caudal end of the ventral hippocampus. Brain sections were immersed in a 1% cresyl violet (CV) acetate solution (Sigma-Aldrich, C5042) for approximately 5 minutes. Subsequently, the slides were immersed in 70% EtOH for 10 minutes. Further removal of excess stain was achieved by immersing the sections in 95% EtOH with 150 ul of acetic acid for approximately 3 minutes, followed by two series of 5-minute baths of 100% EtOH and xylene. Finally, the slices were cover slipped using Eukitt® Quick-hardening mounting medium (03989, Sigma-Aldrich), and imaged under a microscope (Keyence, BZ-X800). The protocol was inspired by [66]–[68]. Slides were visually and quantitatively examined for differences between the control and the KA-treated CA1, CA3 and DG regions using available pig brain atlases. [64], [69].

In each animal, five hippocampal sections were selected along the anterior-posterior axis, spanning the anterior, middle, and posterior hippocampus. Sections were imaged at 40x magnification using a 3x3 stitched configuration and within each section the CA1, CA3 and DG regions were identified using available pig brain atlases. Within one hemisphere, three regions of interest (ROIs) were taken to represent the superior, middle, and inferior parts of each subregion for a total of 9 ROIs per hemisphere per section. Using a standardized 1x1 square centered in each image, neurons were manually counted in QuPath (version 7.0) by a single blinded observer. Neurons were identified by size and a pale nucleus containing a prominent, darkly stained nucleolus. Smaller, darkly stained glial cells with condensed, irregularly shaped nuclei were excluded from counts.

Neuronal counts (𝑋_𝑙𝑒𝑠𝑖𝑜𝑛𝑒_) were Z-score normalized to the non-lesioned hemisphere for each animal and region (CA1, CA3, and DG) using the average (μ_𝑛𝑜𝑛−𝑙𝑒𝑠𝑖𝑜𝑛𝑒𝑑_) and standard deviation (σ_𝑛𝑜𝑛−𝑙𝑒𝑠𝑖𝑜𝑛𝑒𝑑_).

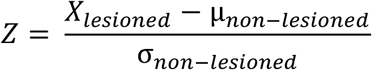

This method accounts for inter-animal variability, regional differences in neuronal counts, and impact of electrode implantation injury. Normalized cell counts were compared using a non-parametric Mann-Whitney U Test at statistical significance of 5%.

## Glossary of Abbreviations

ANT: Anterior Nucleus of Thalamus
AUC: Area Under the Curve
CT: Computed Tomography
CV: Cresyl Violet
DBS: Deep Brain Stimulation
DG: Dentate Gyrus
FBTCS: Focal to Bilateral Tonic-Clonic Seizures
FGATIR: Fast Gray Matter Acquisition T1 Inversion Recovery
HFO: High Frequency Oscillations
HPC: Hippocampus
ICC: Intraclass Correlation Coefficient
IEDs: Interictal Epileptiform-like Discharges
INSR: Implantable Neural Stimulation and Recording
KA: Kainic Acid
LFP: Local Field Potential
MPRAGE: Magnetization-Prepared Rapid Gradient Echo
MRI: Magnetic Resonance Imaging
mTLE: mesial Temporal Lobe Epilepsy
MTT: Mammillothalamic Tract
PPV: Positive Predictive Value
SE: Status Epilepticus
SPEP: Single Pulse Evoked Potentials

## Acknowledgement

Cadence Neuroscience Inc. provided the workstation and implantable hardware. We thank the XRI Core and Department of Comparative Medicine within Mayo Clinic for supporting our experiments. We thank Certicon a.s. for providing CyberPSG and to Wavesurfers s.r.o. for providing EEG Wave that enabled us to review and analyze our data. Figure drawings were made by the manuscript authors. The renderings of the workstation and implantable systems were provided by Cadence Neuroscience.

## Data Availability

Data is available from the corresponding authors upon reasonable request.

## Author Contributions

FM, DM, SC, GW – creating figures, and writing the first draft of manuscript, incorporating suggestions and edits

GW, SC, NG, VKremen, KL – methodology, experimental design, supervision

IK, JVG, KM, JK - surgical protocol development & sterile surgery

FM, IK, JK, ADL, DMontonye, CG - animal care, data recording, charging of the implanted devices

FM, BW, VKrakorova – MRI data processing, image registration, rendering of Papez circuit using histology atlas (Figure 2)

CYK, FM – data annotation and validation of seizure and spike detectors FM, PB, DM – electrophysiology data curation and analysis

GW, NG – electrophysiology data interpretation

DM, SC – histology protocol development, tissue processing and analysis (Figure 4)

ESH, SS, LHK – blinded analysis of histological slices

All authors participated in the manuscript editing.

