## Supplementary Materials for "Kainic Acid Pig Model of Hippocampal Epilepsy"

### Software Tools

Here we provide a complete list of software tools utilized within this project.

#### Data File Management

Multiscale Electrophysiology Format (MEF) as the main format for EEG. MEF is compatible also with the Brain Imaging Data Structure (BIDS). A Python package Meftools was utilized to read and write from and to MEF files.

Repository:
- https://github.com/bnelair/mef_tools

#### Video Recording

Video was acquired using Blue Iris software: https://blueirissoftware.com/

#### EEG Review

EEG Review of MEF files can be performed in either CyberPSG, WaveSurfers or SignalPlant.

Software Links:
- https://cyberpsg.cz/
- https://wavesurfers.science/
- https://www.medisig.com/signalplant/

#### Signal Processing

Publicly available libraries were utilized to process EEG signals and statistical analysis: '*NumPy, SciPy, StatsModels*'. Implementation of frequently utilized functionalities were implemented also in '*BrainMaze: Brain Electrophysiology, Behavior and Dynamics Analysis Toolbox - EEG'* toolbox (*Brainmaze EEG).*

- https://numpy.org/
- https://scipy.org/
- https://www.statsmodels.org/
- https://github.com/bnelair/brainmaze_eeg

#### IED Detection

IED detector first published by Janca (Janca 2013) was previously validated by Sladky in humans and canines (Sladky 2021). The code to the original Matlab implementation is available at: https://github.com/EpiReC-ISARG/IED_detector

A pure Python implementation is available at:
- https://gitlab.com/bbeer_group/development/epycom/epycom

- https://epycom.readthedocs.io/

#### Seizure Detection

The seizure detector was developed and validated on human and canine data by Sladky (Sladky 2021). The implementation utilized also in this work is available at: https://github.com/bnelair/brainmaze-torch.

#### Stereotactic Targeting & MRI Visualization

- **Brainlab** was utilized for stereotactic MRI-guided targeting and pre-operative planning of trajectories (https://www.brainlab.com/).
- **Horos** is a MacOS Based tool for DICOM visualization and storage (https://horosproject.org/).
- **3D Slicer** is a multiplatform tool for processing and visualization supporting DICOM imaging structure (https://www.slicer.org/)

### Stereotactic Frame Photos and Schematics

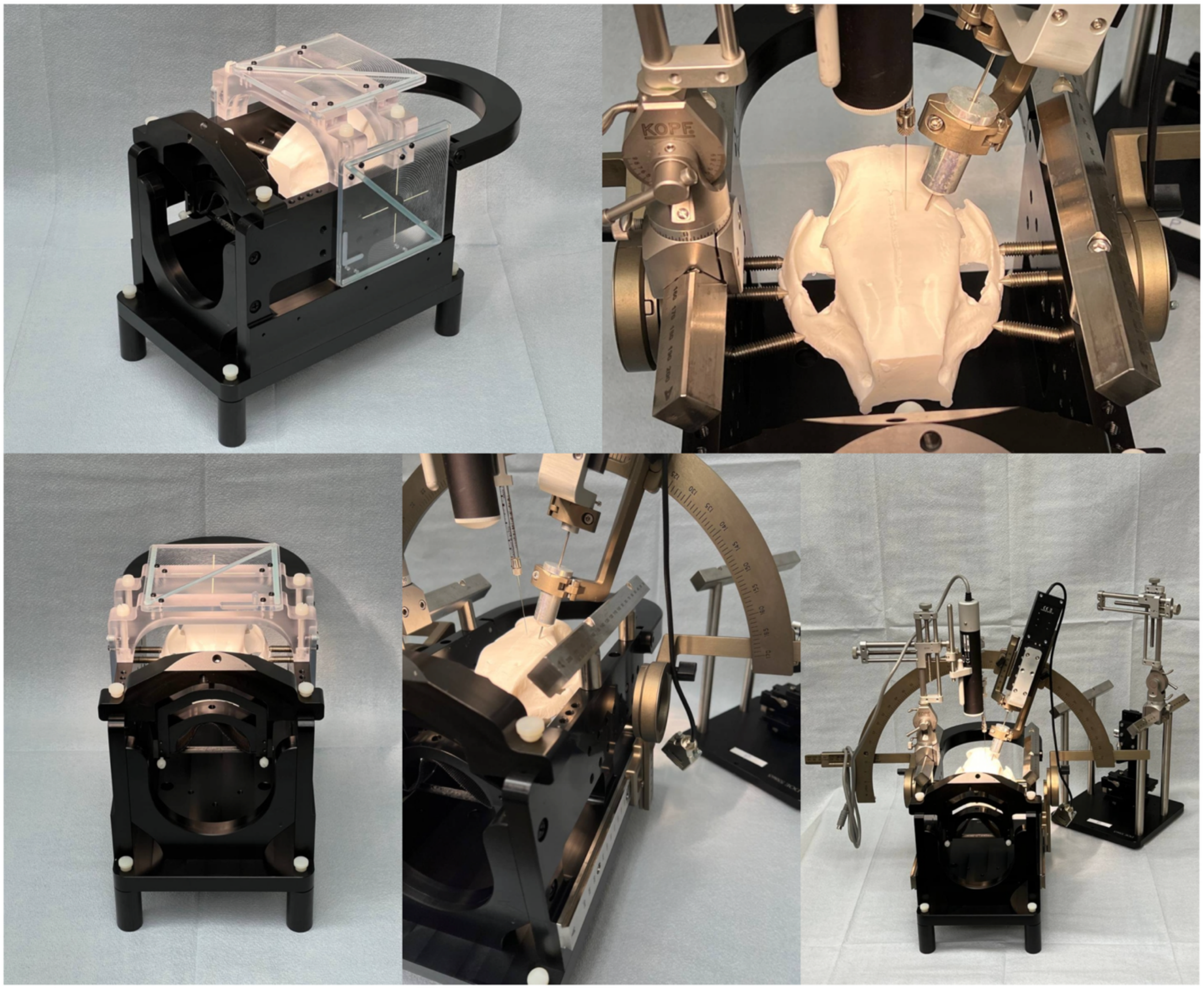

**Supplementary Figure 1.** A photograph of the custom stereotactic frame. The MRI-compatible, machined plastic head fixation frame is compatible with a Leksell arc-frame, Kopf Instruments manipulator and a Hamilton syringe. The frame is utilized for electrode implantation and kainic acid (KA) infusion. The structural design accommodates porcine subjects weighing up to 40 kg.

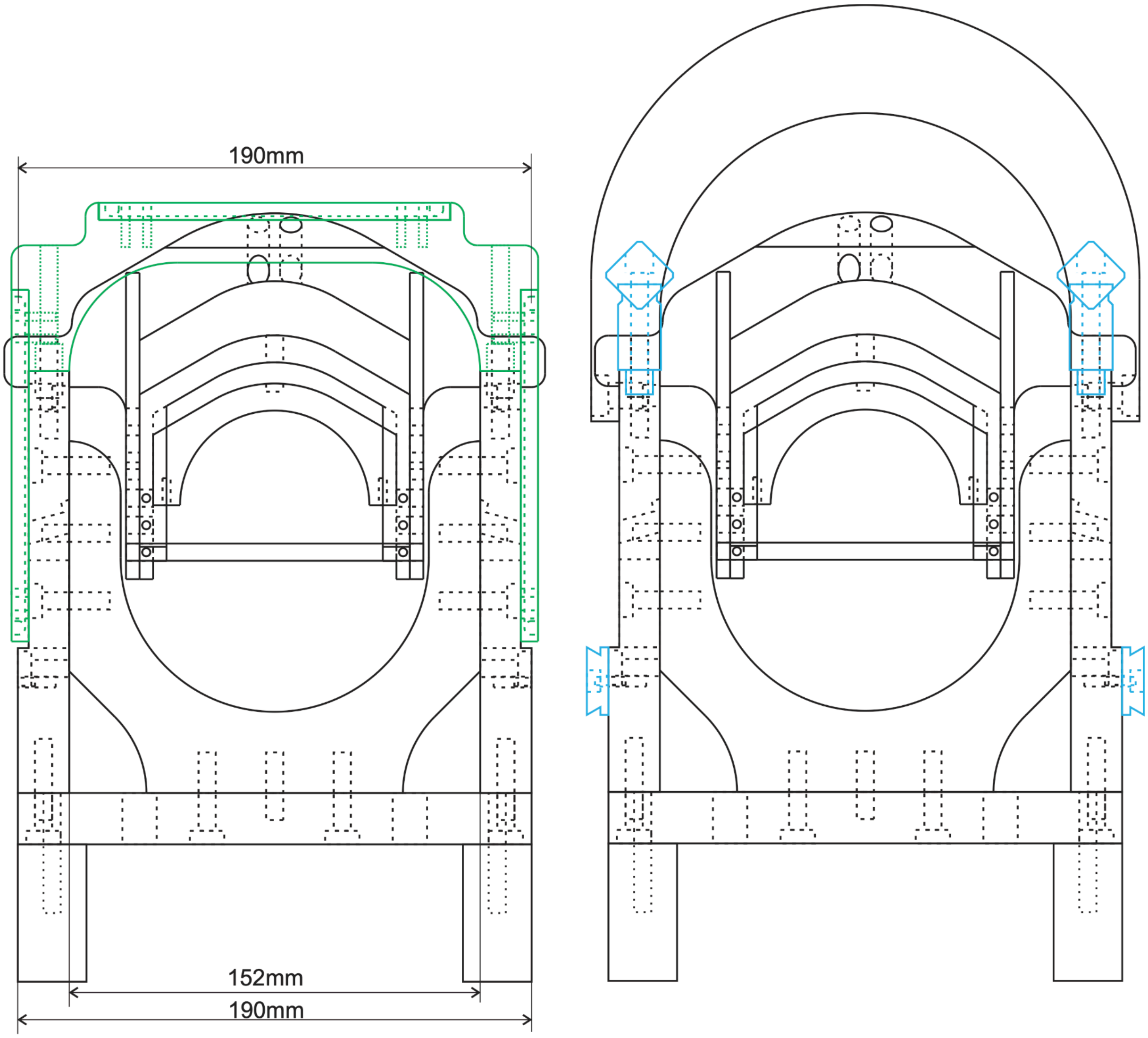

**Supplementary Figure 2.** Coronal schematic drawing of the custom stereotactic head fixation frame. The technical dimensions dictate a maximum exterior width of 190 mm and an interior span of 152 mm.

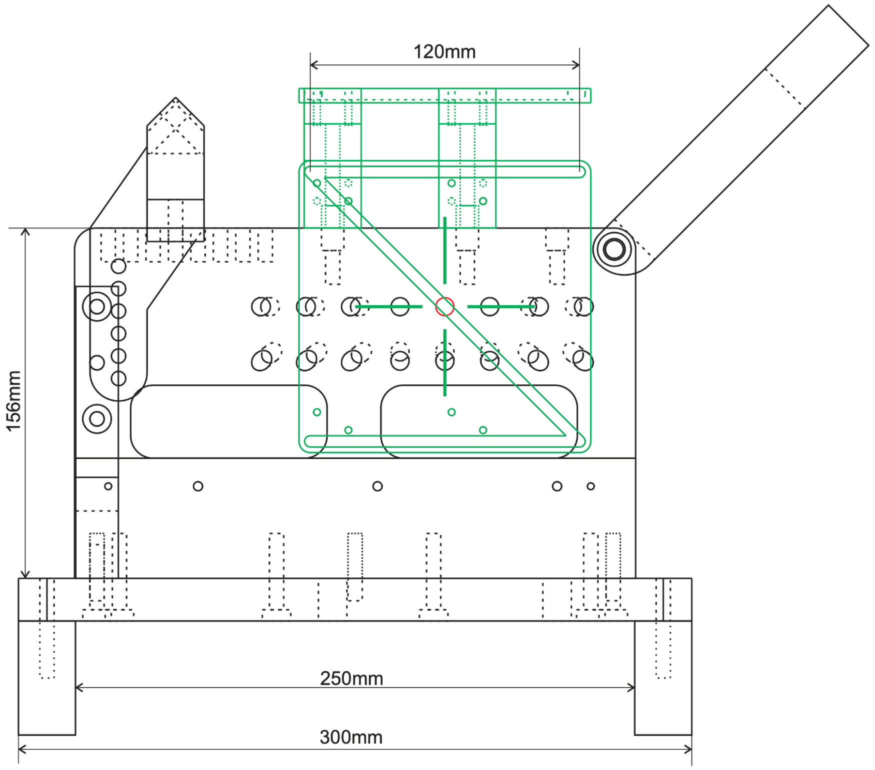

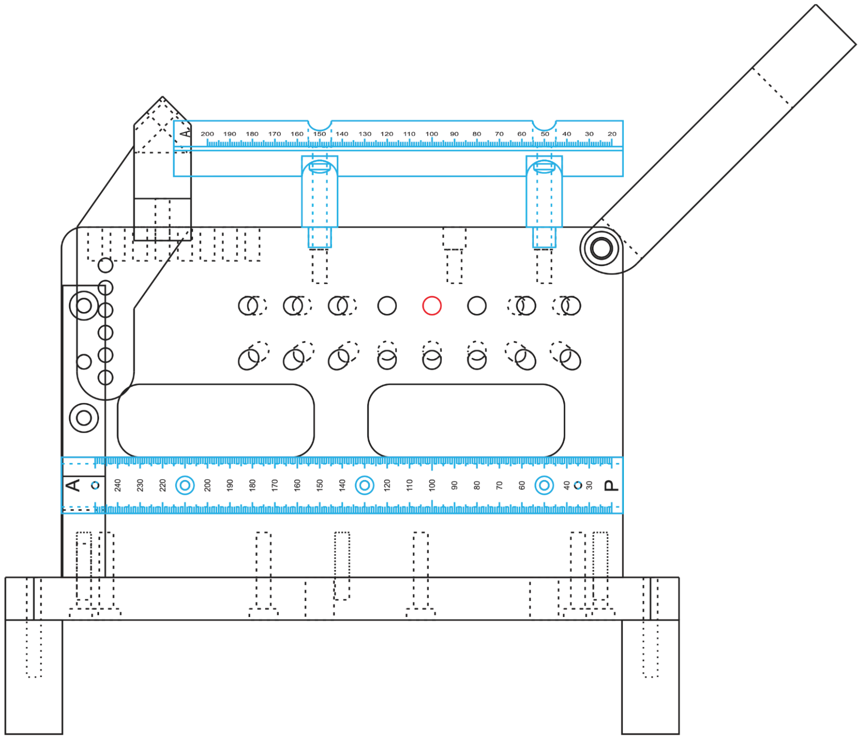

**Supplementary Figure 3.** Sagittal schematic drawing of the custom stereotactic head fixation frame. The specified dimensions document a total base length of 300 mm, an internal base length of 250 mm, and an attached superior component length of 120 mm. The measured height from the base to the specified superior reference point is 156 mm.

### Stereotactic Targeting

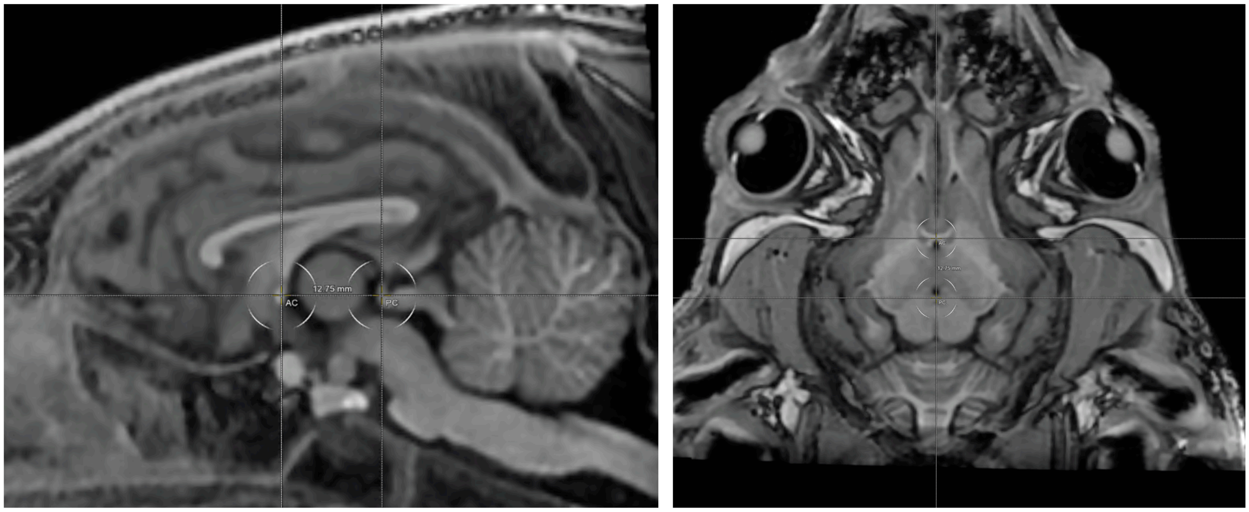

**Supplementary Figure 4. AC-PC Line Alignment and Coordinate System Registration** This figure demonstrates the localization of the anterior commissure (AC) and the posterior commissure (PC), which serve as the primary fiducials for the definition of the AC-PC baseline. T1-weighted Magnetization Prepared Rapid Gradient Echo (MPRAGE) magnetic resonance imaging (MRI) was acquired with isotropic voxel dimensions of 0.8 mm provides spatial resolution necessary for precise anatomical landmark identification. **Sagittal View (Left):** Midline sagittal slice where the AC and PC are identified as hyperintense punctate structures within the third ventricle's boundaries. The AC-PC distance is measured at 12.75 mm, establishing the longitudinal axis of the brain-fixed coordinate system. **Axial View (Right):** Horizontal plane registered to the AC-PC line. The crosshair alignment ensures that the interhemispheric fissure is centered, establishing the mid-sagittal plane for bilateral symmetry and accurate mediolateral and dorsoventral targeting.

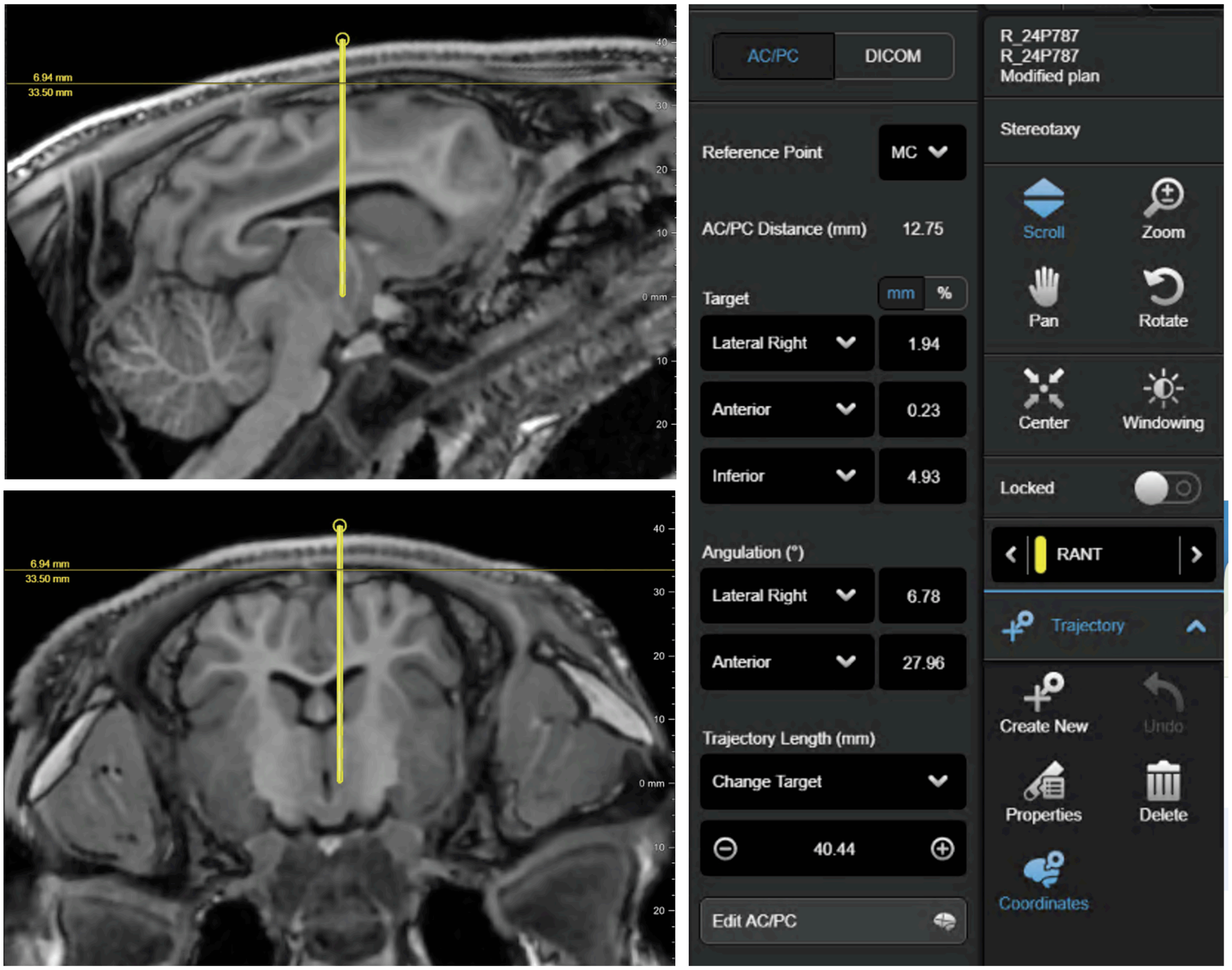

**Supplementary Figure 5. Targeting of Right Anterior Nucleus of the Thalamus (ANT) with AC-PC coordinates**. The Brainlab interface shows a representative targeting of right ANT. **(Left)** The sagittal (top) and coronal (bottom) views display the projected electrode path (yellow line). **(Right):** The provided coordinates are referenced to the mid-commissure (MC – 0,0,0). The target location is calculated based on its spatial offset: Lateral (Right): 1.94 mm, Anterior: 0.23 mm, Inferior: 4.93 mm. The trajectory angulation was set laterally to 6.78° and anteriorly to 27.96°. The trajectory length from the entry point to the top of the skull was identified to 33.5 mm. Trajectory length indicates the length of the whole yellow trajectory end-to-end.

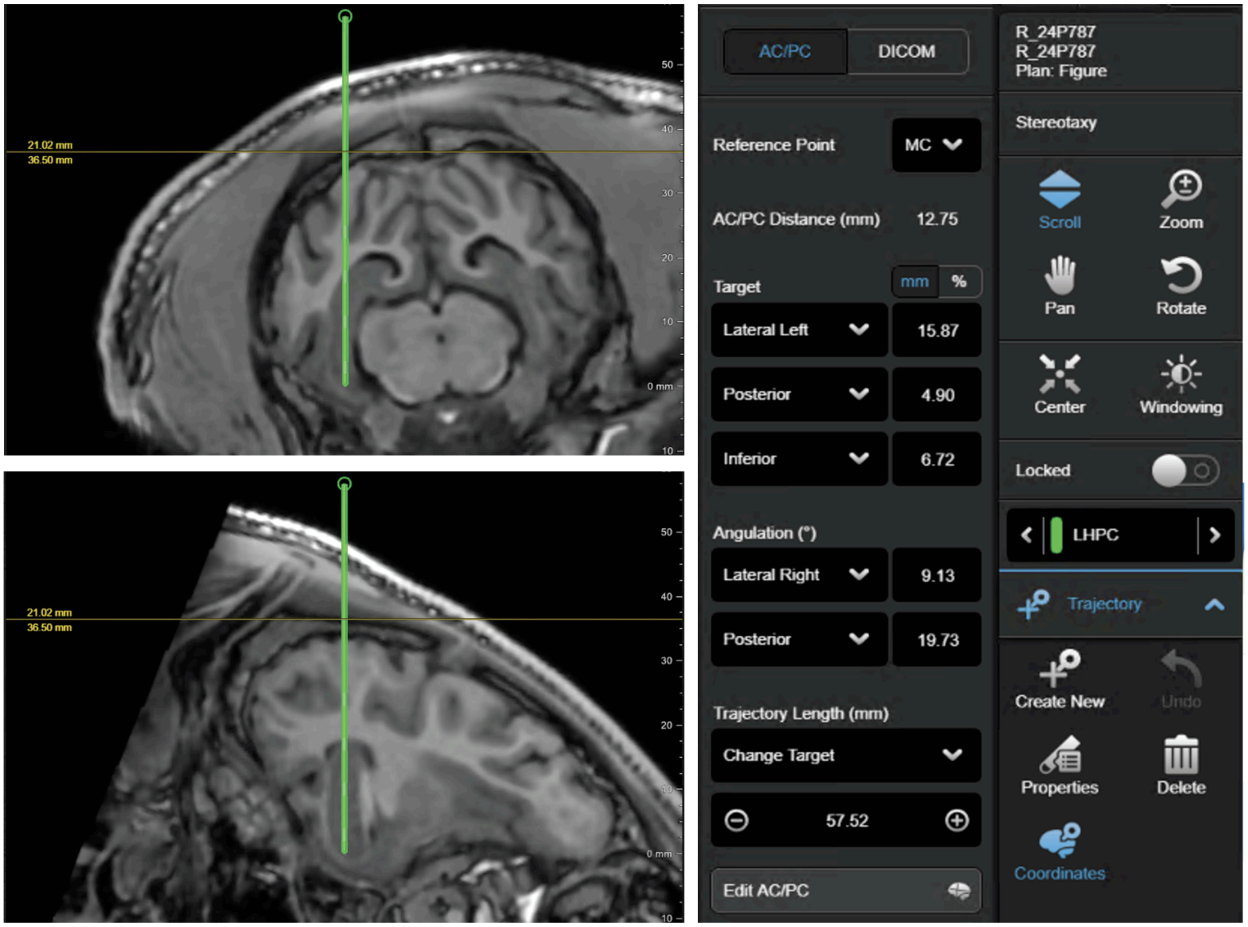

**Supplementary Figure 6. Targeting of Hippocampus (HPC) with AC-PC coordinates**. The Brainlab interface shows a representative targeting of left HPC. **(Left)** The sagittal (top) and coronal (bottom) views display the projected electrode path (green line). **(Right):** The provided coordinates are referenced to the mid-commissure (MC – 0,0,0). The target location is calculated based on its spatial offset: Lateral (Left): 15.87 mm, Posterior: 4.90 mm, Inferior: 6.72 mm. The trajectory angulation was set laterally to the right 9.13° and posteriorly to 19.73°. The trajectory length from the entry point to the top of the skull was identified to 36.50 mm. Trajectory length indicates the length of the whole green trajectory end-to-end.

### Detected Seizure Examples

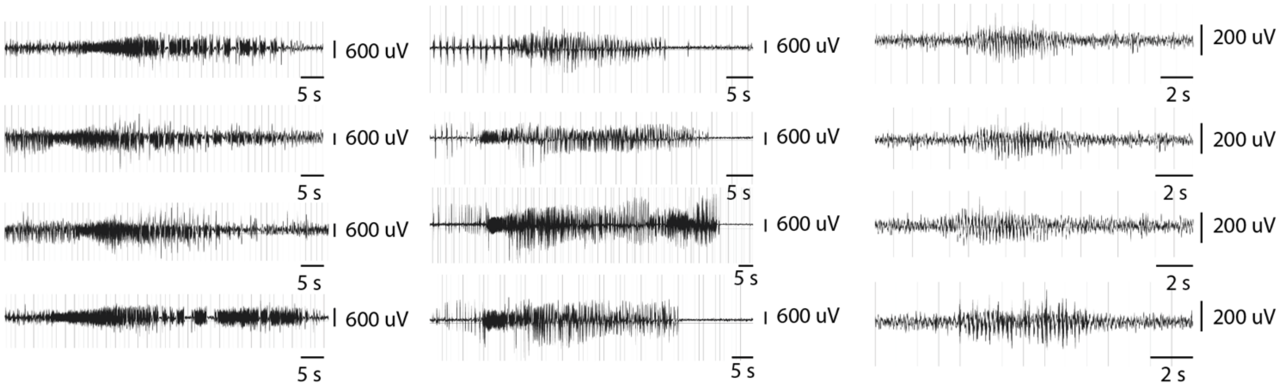

**Supplementary Figure 7. Representative Hippocampal Seizure Trace Examples**. Example LFP traces from 3 KA-treated pigs. Each trace is a single occurrence of electrographic seizure event from an LFP channel at which the detection was performed.
